# Identification and Inhibition of GLUT like proteins in *Trichuris spp* as a druggable target

**DOI:** 10.64898/2026.06.30.735466

**Authors:** Macaulay Turner, Jakub Palinski, Kathryn J Else, Katie Moore

## Abstract

Over a quarter of the world’s population is at risk of infection by soil transmitted helminths (STH). Among the STHs *Trichuris trichiura* infects approximately 7% of people globally, causing a loss of 232,000 DALYS. The main strategy to combat *T. trichiura* infection focusses on mass drug administration with the benzimidazoles. Whilst albendazole and mebendazole have been effective at reducing the burden of other STHs, the cure rate for whipworm is less than 50% with resistance alleles rising. Glucose is the most studied nutrient in *Trichuris spp*, however we have no understanding, at the molecular level of the mechanism of uptake in *Trichuris spp*. We sought to identify putative glucose transporters in *Trichuris* and investigate how these can be inhibited with phloretin. Using the *C. elegans* Facilitated Glucose Transporter 1 (FGT) sequence we identified two potential homologs in *T. muris* (TmGLT) and *T. trichiura* (TtGLT). We should both proteins contained sequence similarity to FGT1 and contained multiple sequence domains associated with glucose and sugar transport. Further, using Alphafold and molecular docking we show glucose docking sites consistent with transport. To asses the ability of phloretin to inhibit glucose transport, we also performed molecular docking with phloretin, showing possible inhibition. To validate the potential inhibition *in vitro* we measured the 48h LC50 of phloretin which we showed to be 111 ug/ml against adult *T. muris* worms, around half that of mebendazole in the same conditions. In contrast phloretin exhibited no effect on worm burden or fecundity *in vivo*.

Together these findings provide the first *in silico* characterisation of putative glucose transporters in *Trichuris spp* and have identified glucose transport inhibition as a promising avenue for anthelminthic drug discovery. Whilst further work is required to optimise *in vivo* efficacy, our results highlight parasite glucose acquisition pathways as potential druggable targets in whipworm.

**Author Summary:** Over a quarter of the world’s population is at risk of infection with soil-transmitted parasitic worms. One of these parasites, *Trichuris trichiura* (whipworm), infects millions of people worldwide. Current treatments rely on two drugs, albendazole and mebendazole, but these are much less effective against whipworm than other parasites, and there is growing concern about resistance. This highlights the need for new treatment strategies.

In this study, we investigated how whipworms take up glucose, an essential energy source required for survival. Using computational approaches, we identified candidate glucose transporter proteins in both the human parasite (*T. trichiura*) and a laboratory model species (*T. muris*). We then tested whether these proteins could be targeted using the compound phloretin. Our results suggest that phloretin may block glucose uptake at the molecular level, and we show that it reduces worm survival in laboratory experiments. However, it did not have the same effect in animal infections.

Together, our findings identify glucose uptake as a potential weakness in whipworms and provide a starting point for developing new treatments.

## Introduction

Over a quarter of the world’s population is at risk of infection by soil transmitted helminths (STH) (1). STH are a major public health burden and collectively cause a loss of 1.9M Disability adjusted life years (DALY) (2).

*Trichuris spp* are STH that infect man and his domesticated mammals with the human infective *T. trichiura* infecting approximately 7% of people globally, causing a loss of 232,000 DALYS (3). *T. trichiura* infection is a disease of poverty, with infection occurring through ingestion of embryonated eggs. Eggs pass through the gut and hatch in the large intestine where larvae go through four moulting stages to reach adulthood (4). Adult parasites inhabit a unique partially intracellular niche, with the anterior head end embedded within the host epithelial cells, and the posterior tail end lying free in the lumen. Adults mate sexually to produce eggs, with a single female worm producing up to 8000 eggs per day, which are passed in the faeces (5).

The main strategy to combat *T. trichiura* infection focusses on mass drug administration (MDA) programmes. School aged and preschool aged children are treated with benzimidazole drugs. Whilst albendazole and mebendazole have been effective at reducing the burden of other STHs, the cure rate for whipworm is less than 50% (6). Benzimidazoles bind the colchicine domain of beta tubulin, preventing depolymerisation. Loss of beta tubulin function results in parasite death by reduction of glucose uptake, and disruption of cellular transport of nutrients. Pertinent to drugs which target gut dwelling nematodes, benzimidazoles are poorly bioavailable, showing low absorption in the digestive tract. Thus, the drug remains primarily in the intestinal lumen where the parasite dwells. Drug resistance is however a major concern, and MDA may be driving similar selection pressure for mutations that has been widely reported in the veterinary field (7, 8). Resistance occurs through polymorphisms in the ITS2 region of the beta tubulin gene (9). Worryingly, a new species of human infective *Trichuris*, *T. incognita* has recently been reported to be highly prevalent and carries benzimidazole resistance polymorphisms (10, 11). Thus, anthelminthics that have an alternative target to the benzimidazoles are urgently needed.

Glucose is the most studied nutrient in *Trichuris spp*. When worms are incubated in glucose free media, their motility is drastically reduced after 17 hours (12) . Further, fluorescently labelled glucose localises within the bacillary band pores, and is possibly taken up into the worm by the bacillary cells that form the bacillary band (12). Although we know glucose is a key nutrient for whipworm, and its likely site of uptake, we have no understanding, at the molecular level of the mechanism of uptake in *Trichuris spp*. The *Trichuris* genome is sparsely annotated and contains no studied glucose transporters. When investigating the proteome of STHs it is often useful to interrogate the proteome of the free-living nematode *Caenorhabditis elegans*. As a model free-living nematode, *C. elegans* has been extensively studied and characterised, used to investigate a plethora of topics from ageing, (13) developmental biology (14) to drug discovery (15). Feng et al identified glucose transporters within the C. elegans genome using BLAST searches. They identified multiple candidates, but showed Facilitated Glucose Transporter 1 (FGT1) was the major transporter, with knockdown of FGT1 resulting in significantly reduced glucose uptake and worm lifespan (16).

FGT1 is a homolog of the human facilitated glucose transporter GLUT1, a transmembrane glucose transporter which is part of the major facilitator superfamily. GLUT proteins have 12 transmembrane alpha helices with a central cavity. Glucose enters the open extracellular confirmation and binds the central cavity, GLUT1 undergoes a confirmational change to its intracellular pose, which releases glucose into the cell (17).

Phloretin is a naturally occurring dihydrochalcone that inhibits FGT1 and other GLUT family glucose transporters. FGT1, shows a Michaelis constant (K) of 2.8 mM for glucose. (16, 18). Despite phloretin’s nonspecific inhibition of mammalian GLUT transporters it can be administered safely to mouse and rat models of a variety of diseases without ill effect (19–21).

Additionally, phloretin has been shown to positively modulate the microbiome, reduce inflammation and increase barrier function in the colon of mice in a colitis-associated colorectal cancer model (22). Further, inhibition of glucose uptake by phloretin has been shown to be directly cytotoxic to cancer cells, with positive effects in breast cancer (23). Although cancer cells and parasitic worms differ profoundly in their physiology both share convergent pressures within their host of effectively acquiring glucose. Where cancer cells require increased GLUT expression to increase their glucose uptake, *Trichuris* must effectively scavenge glucose from their epithelial cell niche. This creates an analogous functional dependency on efficient glucose uptake in their host, disruption of which could be a novel target in *Trichuris* infection. Therefore, as phloretin can disproportionately target cancerous cells, it may also disproportionally affect *Trichuris* over the host gut epithelial cells.

Here, we use the FGT1 sequence to identify a GLUT like receptor in *T. muris* and *T. trichiura*. Using bioinformatic tools and modelling we explore and characterise *in silico* the *Trichuris* GLUT homologs including how they can be inhibited. Finally, we confirm mRNA expression in in *T. muris* and explore the anthelminthic activity of phloretin against whipworm in vitro and in vivo.

## Methods

### Bioinformatic pipeline

The *Caenorhabditis elegans* FGT-1 amino acid sequence (O44827) was retrieved from WormBase and used as a query in BLASTp searches against *Trichuris* genomes hosted on WormBase ParaSite (Version WBPS19). BLASTp searches were conducted using default parameters with the top sequence for *T. trichiura* and T. *muris* selected for further work (24).

Protein sequences identified were aligned using CLUSTAL Omega 1.2.4 with default gap penalties (25). Percentage identity and similarity were calculated from pairwise alignments using the BLOSUM62 substitution matrix.

Functional domain annotation was performed by submitting protein sequences to InterPro 108.0 (25). Annotated functional domains associated with transmembrane transport were retained for downstream analysis.

Protein topology was predicted using TopCONS (26) and Orientations of Proteins in Membranes (OPM (27)), using default parameters. Predicted transmembrane regions and membrane orientation were recorded.

Three-dimensional structural models were generated using AlphaFold v3 (28) Model confidence was evaluated using predicted Local Distance Difference Test (pLDDT) scores, and low-confidence regions were recorded for interpretation of downstream docking studies. Structural similarity between models was assessed using TM-align (29).

Molecular docking was performed using HDock (30) with AlphaFold-predicted protein structures as receptors. Glucose (PubChem CID 5793) and phloretin (CID 4788) ligand structures were obtained from PubChem. Docking was performed under default conditions, and the highest-scoring poses were selected for analysis. For FGT1, and TmGLT undirected docking was performed, for TtGLT with phloretin, docking was directed to the transmembrane domains.

Structural visualisation and figure generation were performed using ChimeraX v1.11.1

### Animal work

All mice used were severe combined immunodeficient (SCID) mice, provided ‘in-house’ by the Biological Services Facility at the University of Manchester. All mice used were male aged between 6 and 8 weeks. Mice were infected with 150 (*in vitro* worm generation) or 20 eggs (*in vivo* efficacy). All animal work was approved by the University of Manchester Animal Welfare and Ethical review Board and performed under the regulation of the Home Office Scientific Procedures Act (1986). Experiments were performed under Home Office License PP3839006 and conbfromed to the ARRIVE guidelines.

### *In vitro* analysis of phloretin efficacy

SCID mice were infected with 150 *T. muris* eggs, at day 35 post infection, mice were euthanised, ceca and proximal colons removed, and worms removed gently using fine forceps. 5 male and 5 female *T. muris* worms were placed in 500 ul of RPMI (Sigma Aldrich) media with 5 x penicillin streptomycin (Sigma Aldrich) per exposure. Phloretin (Thermo Fischer) was dissolved in DMSO (Sigma Aldrich) then added to each exposure group to a final concentration of: 200, 150, 100, 75, 50, 25, 10, 5, 1, 0 ug/ml with a maximum DMSO concentration of 5% volume. The 0 ug/ml control contained 5% DMSO as a vehicle control. Worms were then incubated at 37 °C 5% CO_2_.

Motility was measured at 4h, 24h, and 48h using a motility score of 0 – 3, 0 = no movement in 10 seconds, 1 = anterior end moving only, 2 = slow whole body movement, 3 = normal motility (31). Lethal concentration 50 (Lc50) was calculated using GraphPhad prism using a nonlinear fit model.

### *In vivo* analysis of phloretin efficacy

Mice experienced no adverse effects after oral dosing with phloretin up to a dose of 200mg/kg (data not shown). To evaluate the efficacy of phloretin in worm clearance mice were infected with 20 *T. muris* eggs. After 35 days mice were split into 3 groups at random, group 1 received a single dose of 200 mg/kg phloretin at day 35 post infection, group 2 received one dose of phloretin daily from day 35 to day 39 post infection (total of 5 doses), and group 3 received 5% DMSO contained in sterile water daily from day 35 to day 39. Faecal pellets were collected at day 36, day 38 and day 41 post infection. Faecal pellets were weighed, then eggs isolated via salt floatation. Faecal pellets were weighed then added to 4 ml saturated NaCl, pellets were broken up then left for 1h at R.T. to ensure thorough homogenisation. Suspensions were vortexed and left again for 2h, 800 ul was removed from the top of the suspension and added to a McMaster chamber, eggs were then counted and converted to eggs per gram of faeces. At day 42 post infection, mice were culled and worm burdens measured.

## Results

### Sequence analysis of *Trichuris* FGT1 homologs

There are no characterised glucose transporters in *Trichuris* species’ genomes. FGT1 is the major glucose transporter in the free-living nematode *C. elegans*. To identify the presence of a GLUT-like protein in the *T. muris* and *T. trichiura* proteome, the amino acid sequence of FGT1 (O44827) was used in a BLASTp search. Two homologs were identified, the *T. muris* protein A0A5S6Q2T5 (hereafter referred to as *T. muris* GLUT like transporter (TmGLT)), and the *T. trichiura* protein A0A077Z5C9 (TtGLT). TmGLT and TtGLT had low E-Values, 1.7 E - 65, and 5.4 E-38 respectively and were similar length proteins (FGT1 510 amino acids, TmGLT 490 amino acids and TtGLT 518 amino acids).

Pairwise alignments of FGT1, TmGLT, and TtGLT were generated using CLUSTAL omega (Supporting data Fig 1). Percentage identity and percentage similarities were measured (Table 1) from the pairwise alignments. FGT1 shared ∼40% identity and ∼60% similarity with both *Trichuris* proteins which may indicate relatedness. The two whipworm proteins were more closely related.

**Table 1.** Percentage identity and percentage similarity of FGT1, TmGLT, and TtGLT. Percentage identity and similarity were calculated from CLUSTAL omega pairwise alignments, with a python script used to calculate identity, and similarity based off a BLOSUM62 substitution matrix (25).

| Alignment | Percentage identity | Percentage similarity |
| --- | --- | --- |
| FGT1 vs TmGLT | 44.38 | 62.21 |
| FGT1 vs TtGLT | 43.15 | 60.60 |
| TmGLT vs TtGLT | 84.36 | 88.22 |

Overall percentage sequence identity and similarity were moderate; however no functional information can be inferred from sequence alone. Therefore, InterPro was used to identify predicted functional domains within FGT1, TmGLT and TtGLT. Multiple transporter domains were identified, with FGT1, TmGLT, and TtGLT having 14, 12, and 13 domains respectively that were predicted to be involved in transport, including sugar transport. InterPro analyses indicated a functional relationship between all three proteins. To evaluate whether this relationship reflects homology, we carried out a three-way sequence alignment. Percentage identity and similarity was quantified within the domains identified (Supporting Table 1)). All three proteins contained a major facilitator superfamily (MFS) domain. Given that GLUT transporters are members of the MFS, this domain was selected as a representative region of comparison (Table 2). The percentage identity and similarity between FGT1 and the two *Trichuris* proteins were higher in the MFS domain than the rest of the protein indicating functional homology. Furthermore, over 80% identity between the *Trichuris* proteins paired with shared domain architecture indicates they are orthologs.

**Table 2.** Percentage identity and similarity of the major facilitator superfamily domains identified in each protein using InterPro. Alignments produced using CLUSTAL Omega (25) and percentage identity and percentage similarity calculated using a BLOSUM62 substitution matrix.

| Protein | Start | End | Percent Identity to FGT1 | Percent similarity to FGT1 | Percent Identity with TmGLT | Percent Similarity with TmGLT | Percent Identity with TtGLT | Percent Similarity with TtGLT |
| --- | --- | --- | --- | --- | --- | --- | --- | --- |
| FGT1 | 49 | 495 | - | - | 49.9 | 68.3 | 49.4 | 68.0 |
| TmGLT | 30 | 472 | 50 | 68.2 | - | - | 91.9 | 95.5 |
| TtGLT | 41 | 500 | 49.5 | 68.2 | 91.9 | 95.5 | - | - |

### Predictive modelling of *Trichuris* GLUT like proteins

InterPro identified sugar transport motifs in all three proteins. To investigate whether the sequence similarity translated into structural similarities AlphaFold was used to predict protein 3D structure (Figure 1). Additionally, TopCONS (26) and OPM (27) was used to predict transmembrane domains and topological orientation. All proteins contained 12 transmembrane alpha helices, with a central pore between these TM helices, consistent with GLUT proteins. FGT1, TmGLT, and TtGLT are predicted to have the same topology with their sequence beginning and ending in the cytoplasm.

**Figure 1.**
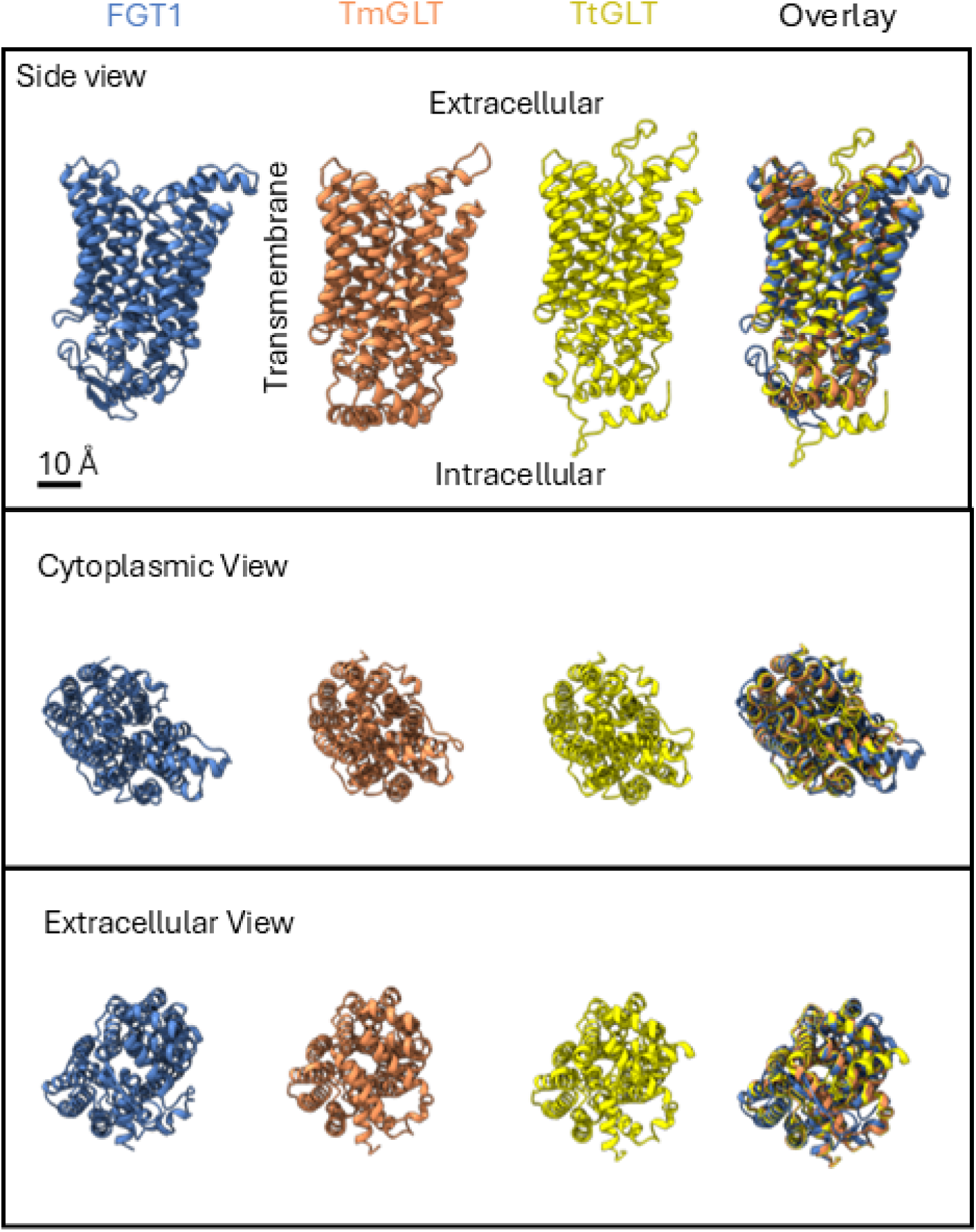
AlphaFold models of FGT1, TmGLT and TtGLT. All three proteins have the same topological orientation calculated via TopCONS and OPM. Figure produced using ChimeraX.

FGT1, TmGLT and TtGLT all appear to be folded in the same orientation and have many overlapping residues. However, to more accurately measure the relatedness of protein structure Tm align was used (29). A Tm score of >0.5 shows relatedness in folding with a Tm score of 1 being identical folding. FGT1 with both TmGLT and TtGLT had a Tm score of 0.89, whereas the two whipworm proteins had a score of 0.97. AlphaFold models of each protein showed near identical folding and secondary structure.

Sequence, and structural analysis implicate FGT1, TmGLT, and TtGLT in glucose transport. FGT1 is the major glucose transporter in *C. elegans* and is a GLUT protein. To characterise the interaction between the proposed substrate glucose, with the FGT1, TmGLT, and TtGLT a docking study was performed using Hdock (30). Figure 2. Shows a representative glucose binding pocket for each protein with the pose with the highest docking score shown.

**Figure 2.**
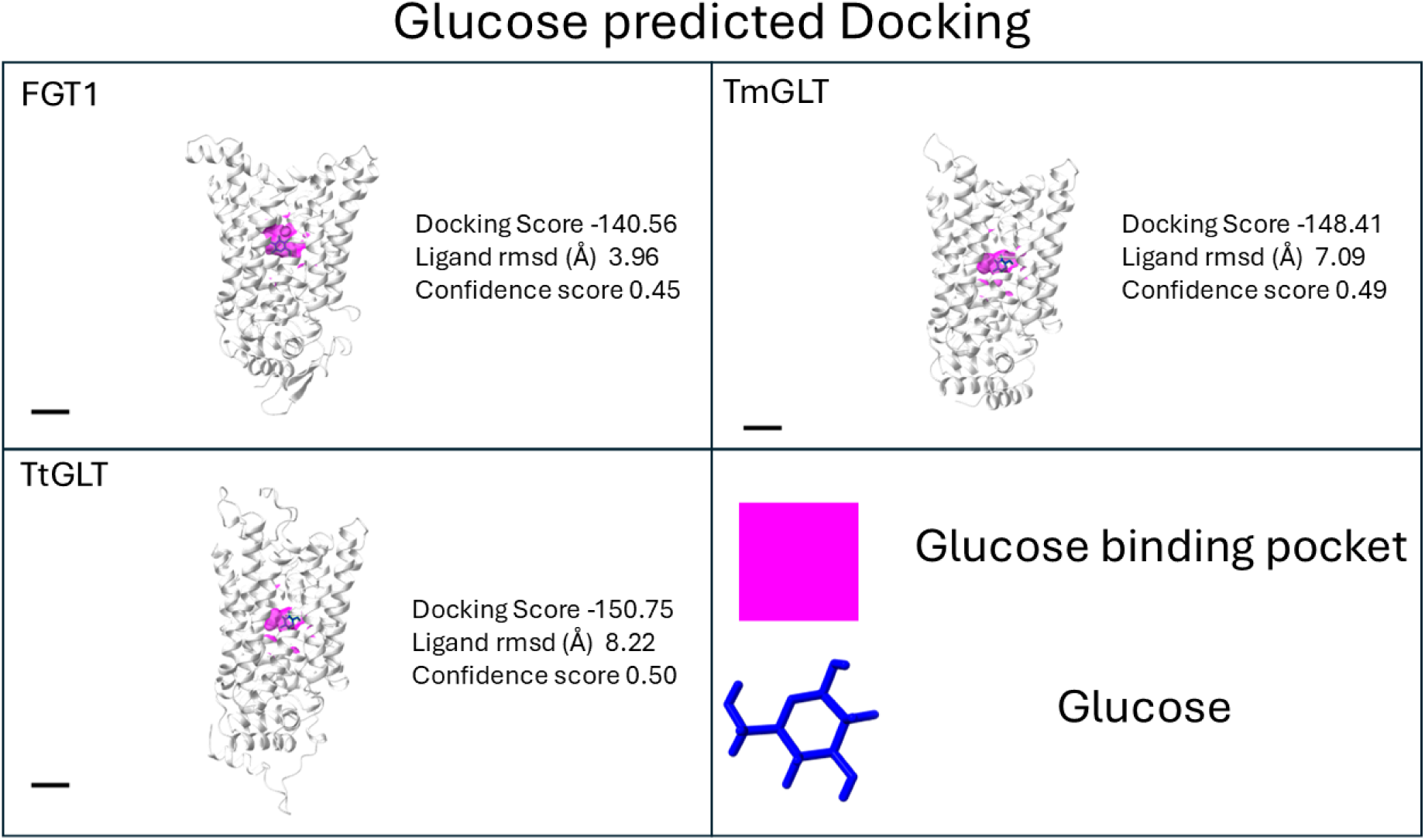
Predicted glucose binding pockets in FGT1, TmGLT, and TtGLT using HDock. All three proteins possess glucose binding pockets within the central cavity of each protein. Docking scores, ligand RMSD, and confidence scores are presented which were calculated using HDock. Figure was produced using ChimeraX. Scale bar = 10 Å.

The highest predicted binding of glucose was within the central cavity of each protein, consistent with GLUT transporter activity. The Docking score shows how strongly the binding of glucose is predicted to be with the protein with a higher negative score being stronger binding. All three proteins were predicted to have a score between -140 and -150. Moderate docking scores shows moderate affinity, which is consistent with transporter activity, rather than strong permanent binding. Glucose binding with GLUT transporters is flexible with flexibility arising from glucose forming multiple weak rotatable interactions that accommodate conformational change of the protein to allow transport across the membrane.

The ligand RMSD values ranged from ∼4 Å (FGT1) - ∼8 Å (TtGLT) reflecting the ability of glucose to adopt multiple orientations within the binding cavity. Adopting multiple confirmational orientations is consistent with GLUT transporter activity. Confidence scores were 0.45 - 0.5 which is defined as ‘possible binding’. Interestingly, FGT1 had the lowest confidence score of all proteins analysed, despite being the only protein confirmed to transport glucose biochemically.

Phloretin is a known inhibitor of FGT1 and GLUT transporters and has been used in murine cancer models to alter progression of disease (20, 23). Thus, inhibition of FGT1 and GLUT transporters is well established both experimentally and using *in silico* approaches. HDock was also used to analyse how phloretin binds to FGT1, TmGLT, and TtGLT (Figure 3). Phloretin was predicted to bind the central cavity of FGT1 and TmGLT with an orientation parallel with the transmembrane alpha helices. However, surprisingly, for TtGLT, phloretin was predicted to bind outside of the membrane pore, despite 22 out of 23 of the binding pocket residues identified in TmGLT being conserved in TtGLT. Structural overlay (Figure 1) and TM scores show high structural homology between TmGLT, and TtGLT. However, when docking was restricted to the transmembrane alpha helices (Figure 2) there was an agreement with docking of phloretin with TmGLT. Therefore, the external binding pose of phloretin with TtGLT is likely due to limitations in rigid docking and side-chain geometry rather than loss of binding pocket.

**Figure 3.**
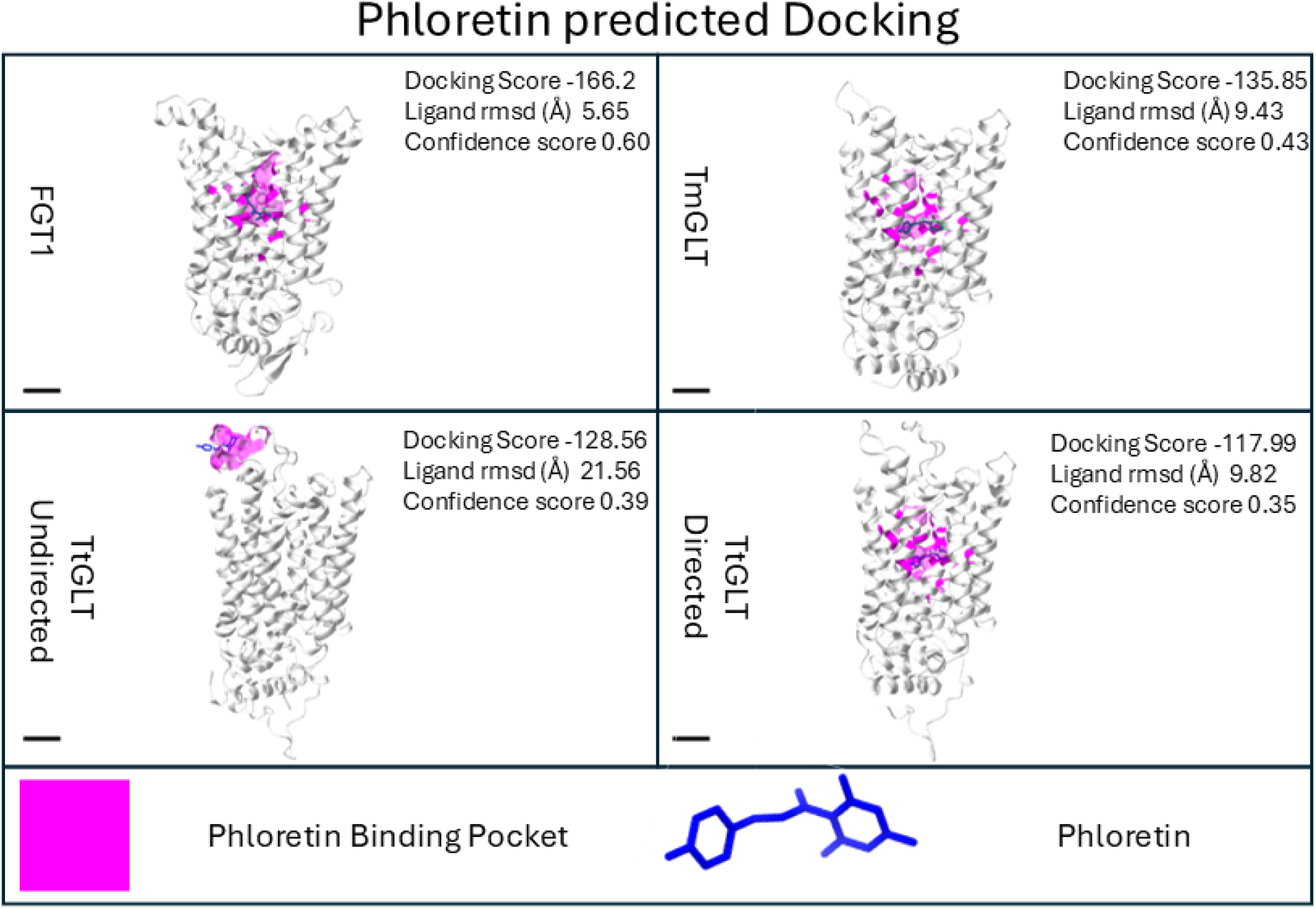
HDock prediction of binding pockets (pink) of phloretin (blue) with FGT1, TmGLT and TtGLT. Docking of phloretin with TtGLT was predicted to be outside the central pore therefore docking was repeated with HDock directed to the transmembrane alpha helices.

Docking scores for phloretin with TmGLT and TtGLT were lower than those obtained for glucose (Figure 2, 3). Despite this, phloretin can still function as an inhibitor. GLUT–glucose interactions are formed through multiple weak, transient interactions distributed along the pore of the transporter, rather than a single high-affinity binding site. As a result, an inhibitor does not need to out-compete the full binding energy of glucose, it only needs to engage the pocket strongly enough to disrupt these transient interactions or physically block substrate passage.

Docking scores, ligand RMSD, and confidence scores are presented which were calculated using HDock. Figure was produced using ChimeraX. Scale bar = 10 Å.

Table 2 shows the amino acid residues that interact with phloretin in the central cavity. There is good agreement showing phloretin is closely associated with transmembrane alpha helices 5 7 and 11. Additionally, phloretin was shown to interact with many of the same residues on the transmembrane helices. Despite modest overall sequence homology between FGT1 and TmGLT or TtGLT the residues predicted to be involved in phloretin binding are conserved.

**Table 2.** Transmembrane alpha helices of FGT1, TmGLT and TtGLT that contain amino acids predicted to be in the binding pocket of phloretin. Binding pocket and docking prediction using HDock, and transmembrane alpha helix prediction was performed using OMD.

| Transmembrane alpha helix |  |  |  |  |  |  |  |  |
| --- | --- | --- | --- | --- | --- | --- | --- | --- |
| Protein | TM1 | TM2 | TM4 | TM5 | TM7 | TM8 | TM10 | TM11 |
| FGT1 | PHE60<br>ILE 64 | N/a | Ser<br>171 | GLN195,<br>VAL 198,<br>THR 199,<br>ILE 202 | GLN 315, GLN 316,<br>ILE 320, ASN 321,<br>MET 324, ASN 352 | N/A | PHE418,<br>TRP 427 | ASN 450, TRP<br>451, ASN 454 |
| TmGLT | PHE41,<br>ILE45 | SER<br>86 | N/a | GLN174,<br>VAL 177,<br>THR 178,<br>ILE 181 | GLN295, GLN 296,<br>ILE 300, ASN 301,<br>MET304 | ASN333,<br>THRE<br>337 | PHE 395,<br>ALA 396,<br>GLY 400,<br>TRP 404 | VAL 424, ASN<br>427, TRP 428,<br>GLY 429, ALA<br>431 |
| TtGLT | GLY 52 | N/a | N/a | GLN 201,<br>ILE 208 | ASN 327, GLN 321,<br>GLN 322, ILE 326,<br>MET 330 | ASN359 | N/A | ALA 456, ASN<br>452, GLY 454, |

### Evaluation of Phloretin as an anthelminthic

We first sought to evaluate the expression of TmGLT in *T. muris*. We therefore analysed the Foth et al RNA seq data set to find the abundance of TmGLT transcripts in the anterior and posterior of worms (32). We found confirmed that TmGLT is expressed throughout life cycles stages and within the anterior and posterior of the worm.

Given that *T. muris* adult stage parasites express TmGLT, and its reasonable percentage identity, glucose transport prediction and structural similarities with FGT1, phloretin was evaluated as an anthelminthic. Adult *T. muris* worms were exposed to 0 – 200 µg/ml Phloretin and their motility measured on a scale of 1 – 3 (31) at 4, 24 and 48 hours. (Figure 4). Phloretin decreased motility of worms at a concentration greater than 5 µg/ml after 4 hours. After 24h some worms had motility of 0 at concentrations of 100 ug/ml with most worms dead at concentrations >150 ug/ml. At 48h, lethal concentration 50 (LC50) was calculated to be 111 ug/ml.

**Figure 4.**
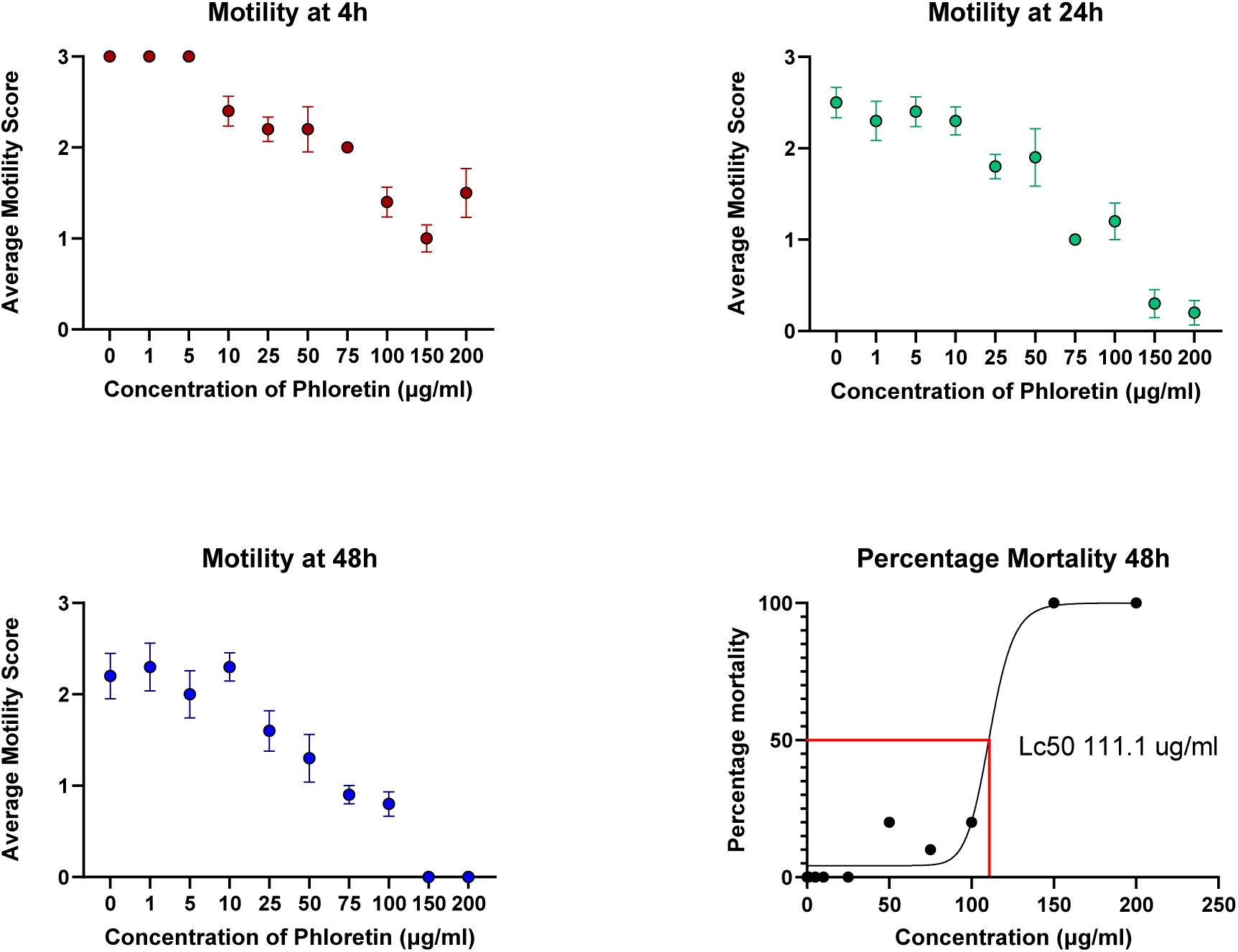
Average motility of 10 *T. muris* worms (+/- standard error mean) exposed to between 0 and 200 ug/ml of phloretin. Individual worms were placed in 500 ul of media with varying concentrations of phloretin. Phloretin was solubilised in DMSO and added with a final DMSO concentration of 5%. Vehicle control was 5% DMSO alone. Worms were incubated at 37 °C 5% CO2. Motility was scored on a scale of 0 – 3, 0 = no movement in 10 seconds, 1 = anterior end moving only, 2 = slow whole-body movement, 3 = normal motility. Motility was measured at 4h, 24h, and 48h (n=10 individual whipworms). LC50 was calculated at 48h. produced using GraphPad prism.

Given that phloretin showed *in vitro* efficacy against adult *T. muris* parasites, we sought to assess the ability of phloretin to reduce worm burden *in vivo*. 6 mice per treatment group were infected with 20 infective *T. muris* eggs and treated by oral gavage at day 35 with: 1 dose of (200 mg/kg) phloretin, 1 dose of phloretin daily for 5 days, or 1 dose of DMSO daily for 5 days. Weight was measured daily after exposure, mice lost minimal weight over the course of treatment regardless of group (data not shown). Lack of weight loss indicated that there was no significant interaction of phloretin with host GLUT transporters.

Faecal pellets were collected at day 36, 38, and 41 days post infection. Eggs/gramme faeces’ (indicative of worm fecundity and thus overall worm health) was not significantly different between DMSO control and either treatment group (Figure 5).

**Figure 5.**
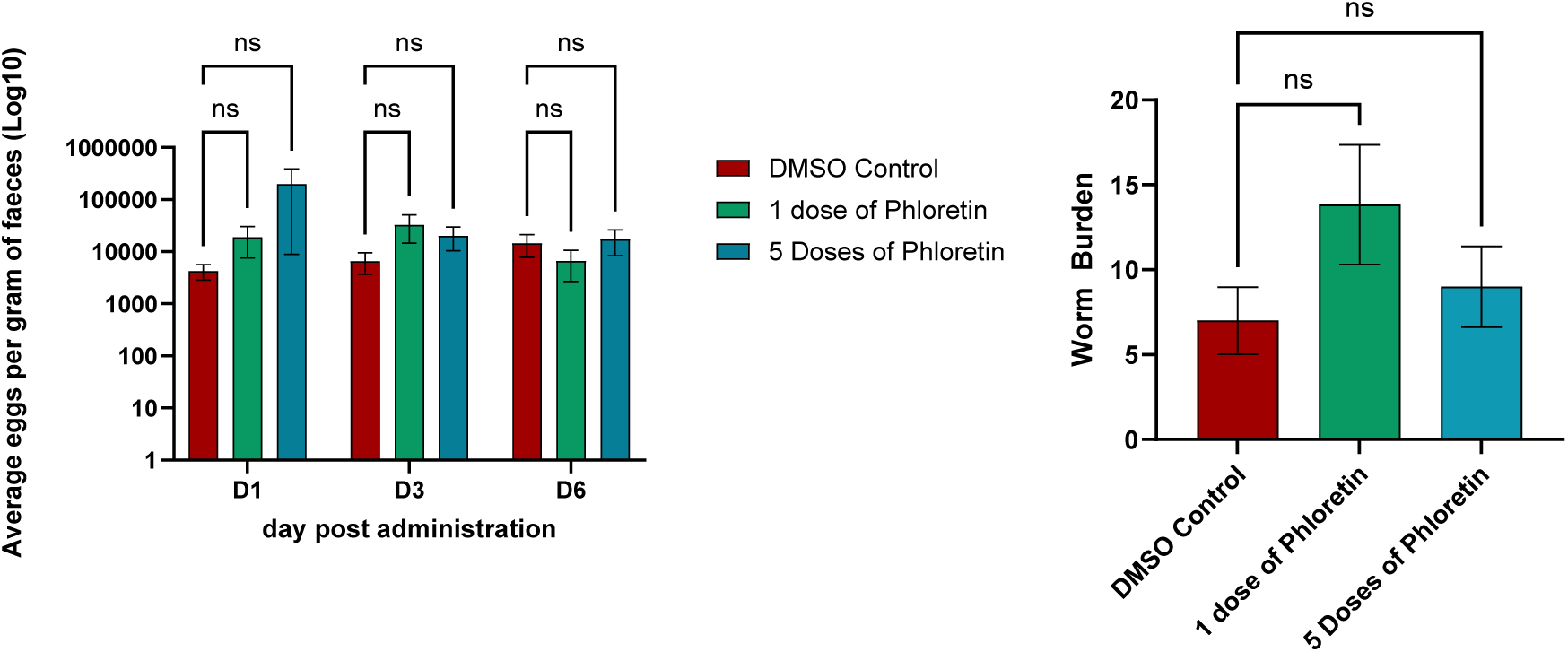
Effect of phloretin on *T.muris* fecundity and worm clearance *in vivo*. 6 SCID mice per group were infected with 20 *T. muris* eggs. At day 35 mice were treated with either 1 dose of phloretin, 1 dose of phloretin daily for 5 days, or 1 dose of 5% DMSO vehicle control daily for 5 days. At day 1, 3 and 6 post administration of phloretin fecael pellets were collected and eggs counted. At day 7 post administration of phloretin, mice were euthanised and worm burden taken. No difference in *T. muris* fecundity or burden was seen. Figure produced using GraphPad Prism, statistical test was a 2 way ANOVA, graph shows +/-standard error mean.

At day 42 post infection mice were culled and worm burdens quantified. There was no significant difference in worm burden between treated and control groups. Therefore, phloretin treatment via oral gavage was not effective in clearing T. muris infection at the dose and administration regime employed in this experiment.

## Discussion

Infection with STH is a major burden on public health in the world’s poorest communities (2). Despite efforts of benzimidazole MDA programmes, infection with STH including *T. trichiura* remains extremely common, with drug resistance being a major concern (6). Therefore, novel anthelminthics with a target different to that of the benzimidazoles is urgently needed.

*Trichuris* has been shown to have reliance on glucose, with worms incubated in glucose free media quickly dying (12). Therefore, we sought to identify and characterise the glucose transporters in *Trichuris*. Further, we aimed to explore the use of targeting these transporters with phloretin, a molecule that has been shown to inhibit GLUT transporters, and has been used successfully in renal and cancer models (19–21, 23)

The BLAST results of the FGT1 gene within the *T. muris* and *T. trichiura* genomes produced TmGLT and TtGLT as potential homologs. Modest percentage identities and similarities between TmGLT, TtGLT and FGT1 were seen, however given the evolutionary distance between *C. elegans* and *Trichuris* this is not surprising. Indeed Feng et al showed FGT1 had a low sequence homology to human GLUT transporters, but was still able to transport glucose (16). Furthermore, the Tm score of 0.89 shows extremely conserved folding between each of the three proteins.

In addition to sequence, domain and folding similarities we also show the predicted docking of glucose with each protein. Glucose is predicted to bind within the central cavity of each protein, consistent with glucose transporter activity. Despite docking predictions showing different orientations of glucose in the transporter, along with different RMSD values between proteins. this was consistent with glucose transporter activity. Glucose enters the outward open confirmation of GLUT transporters then forms multiple weak interactions with amino acids inside the central cavity (17). Therefore, it is unsurprising that HDOCK has predicted multiple different orientations and binding pockets rather than one fixed position.

HDOCK was also used to predict the interaction between phloretin and each protein. The model showed phloretin in a central binding pocket within the central cavity of each protein. HDOCK confidence scores were low for each prediction. Docking scores have known limitations, particularly regarding protein confirmational flexibility. Further, *Trichuris*, and indeed other helminth proteins, have post translational modifications that are difficult to model (33, 34). Additionally, FGT1 has a low confidence score when docked with glucose in HDock. Previous work has shown biochemically that FGT1 is a glucose transporter (16, 18) Therefore, docking should be interpreted as a hypothesis rather than a definitive prediction of binding pocket. Phloretin was also predicted to interact with the central cavity of each protein. Phloretin is a competitive inhibitor of GLUT transporters, entering the outward facing conformation of GLUT transporters and binding preventing alteration to the inward facing conformation (35).

TmGLT is expressed throughout L2, L3 and adult stages of *T. muris* (32). Interestingly it is most highly expressed at L2 stages. *T. muris* undergoes 4 moulting stages increasing drastically in size from ∼80 µm in L1 to ∼3 cm in adults, representing a substantial increase in biomass. Such rapid growth likely requires significant glucose uptake, which aligns with the reported mRNA abundance in L2 vs adults (2.42 vs 2.04). Despite reduced expression in adults, Foth et al showed expression is retained suggesting a continued functional role beyond development. Therefore, we investigated the effect of inhibiting TmGLT in adult parasites using phloretin.

Phloretin showed a dose dependent effect on worm motility *in vitro* , with a low concentration of 10 ug/ml decreasing motility even after 4 hours. Phloretin had an LC50 of 111.1 ug/ml against T. muris after 48 hours, which is considerably lower than the currently used benzimidazoles which have a 72h IC50 of >200 ug/ml (36).

Oral administration of phloretin at a dose of 200mg/kg was well tolerated by mice with no adverse effects, but had no effect on worm burden or fecundity of worms. Thus despite the comparable in vitro LC50 to other efficacious drugs (36) we saw no effect *in vivo*, likely due to insufficient quantity of phloretin being taken up by the worm. Phloretin has low bioavailability when administered orally. It is absorbed most at the proximal colon and cecum, however, is still only reported to have a bioavailability of 8.7% (37), therefore despite favourable pharmacokinetic properties to target the large intestine dwelling whipworm, local intestinal concentrations may still be an issue. As mentioned, the bacillary band has been implicated in glucose transport, with this structure located on the ventral side of the stichosome which is embed within an intracellular epithelial syncytial tunnel (12). If the bacillary band is the site of glucose uptake, it may be the site of TmGLT expression. It is unclear where or how phloretin enters the worm, but if it is acting on the bacillary band, the intracellular niche of whipworm paired with low bioavailability of phloretin may explain the lack of clearance *in vivo*. Low bioavailability may mean phloretin is more efficacious against the fully extracellular STH such as *Ascaris lumbricoides, Nectar americanus* or Ancylostoma *duodenale*. These worms, unlike *Trichuris*, have the full length of their body exposed to the luminal contents, so may be more susceptible to phloretin

The present study was performed using immunodeficient mice, allowing us to identify the role of phloretin on *T. muris* directly, and independently of any potential effect of the adaptive immune response. However, phloretin has been shown to have anti-inflammatory effects in a colitis associated colorectal cancer model, significantly downregulating type 1 cytokines (22). Further phloretin has been shown to directly modulate cytokine release and is implicated in maintenance of epithelial cell tight junction *in vitro* (38). *Trichuris* infection has been well explored in an immune context. Chronic whipworm infection occurs due to a type 1 response, with worm expulsion controlled by type 2 immunity (39). Despite lack of direct effect of phloretin in worm clearance, it is possible that administration of phloretin to *T. muris* infected immunocompetent mice may supress the Th1 response driving a Th2 response and expulsion. Reliance on host immune response in drug treatment has been observed in other parasitic infections, notably in praziquantel treatment of *Schistosoma spp* where praziquantel damages the parasite’s tegument facilitating immune-mediated clearance (40).

In the current study, phloretin was delivered using a DMSO suspension in water. Previous studies have shown that the bioavailability of phloretin can be increased by using mixed polymeric modified self-nano emulsions (SNEs) (41). When phloretin is encapsulated in SNEs its solubility increased 3000-fold and its bioavailability 7-fold (41). Delivery in SNEs was also shown to increase the anti-inflammatory effect of phloretin. Thus, using SNEs in phloretin delivery in a *T. muris* model could increase its efficacy in worm clearance, cautioned by the fact that a higher bioavailability could increase adverse effects, and off target interaction with host GLUT transporters. Employing different delivery methods to alter compound pharmacokinetic properties is a fast-evolving field, which will undoubtedly be implicated in future anthelminthic research.

Despite the lack of efficacy in worm clearance, we have identified a novel target for drug discovery. *Trichuris* worms have a strong dependency on glucose and are quickly killed without access to it. Other small molecules could be used to target *Trichuris* GLUT like transporters, and this is an avenue for future research. Further, different vehicle delivery methods and effects in immunocompetent models present exciting opportunities. The use of phloretin represents a step forward to understanding *Trichuris* biology, and how we can exploit this to reduce the burden of this prevalent infection.

## Acknowledgements

The authors would like to acknowledge Dr Ömer Faruk Bay for his assistance in analysing *T. muris* mRNA seq data set.

## Data availability

All relevant data are within the manuscript

## Author contributions

MJT, KJE and KLM conceived the project. MJT and JP carried out the experiments and analysed the data; MT, KJE and KM wrote the paper. All authors reviewed the manuscript.

## Supporting information

Supporting data Figure 1. Clustal Omega alignment of TmGLT, TtGLT, and FGT1.

Supporting data Table 1. Domains in TmGLT, TtGLT, FGT1 identified by Interpro predicted to be involved in sugar transport and the percentage identity and percentage similarity in these domains. Percentage Identity and percentage similarity were calculated using a BLOSUM62 matrix with pairwise alignments generated using Clustal Omega.

